# Glophyt: a user-friendly, general-purpose program for nonlinear and multidimensional curve fitting via a hybrid stochastic and deterministic approach

**DOI:** 10.1101/2024.04.25.591093

**Authors:** Georges Czaplicki, Serge Mazeres

## Abstract

**Background:** Model validation depends on the agreement between the predicted and experimental data. However, finding solutions to problems, described by equations with many parameters, for which virtually nothing is known, is a difficult task. For example, the extraction of kinetic parameters from complex schemes representing the conversion of a substrate into a product by an enzyme in the presence of an inhibitor is extremely difficult, as even the orders of magnitude of the parameters are not known. This makes curve fitting very difficult in case of multidimensional and nonlinear data. This article presents a graphical user interface-based program employing a hybrid stochastic and deterministic approach, which allows for easy and reliable determination of model parameters.

**Results:** The program has been extensively used in several laboratories at our institute and has proven to be efficient in determining model parameters in many different fields. Although its origins are related to kinetic studies in enzymology, it has been successfully tested on data from various sources, such as pharmacological studies of ligand−receptor binding, entomological studies of populations, bacterial growth, photosynthesis, toxicology, differential scanning calorimetry, isothermal titration calorimetry and nuclear magnetic resonance spectroscopy.

**Conclusions:** This program presents an effective solution for researchers facing the problem of extracting model parameters from multidimensional and nonlinear data where even the orders of magnitude of parameters are not known. Its graphical user interface makes it easy to use, does not require any programming skills, and it is cost-free. It is available for Windows and Linux platforms.

## Background

It is a common practice in experimental science to test different models of the studied phenomena by comparing measured results with those calculated on the basis of a given equation. The purpose is to extract the values of inherent parameters to gain a better understanding of the underlying processes. A good example of such a situation is provided by kinetic studies of enzyme-inhibitor reactions, which require time-consuming analyses, where fitting various conceptual models of interaction pathways to experimental data plays an important role. To find a set of kinetic constants consistent with the observed data, nonlinear multidimensional global optimization techniques must be used (see (1) and references therein). There are a number of software solutions available to accomplish this task, which minimize differences between the predicted and experimental curves. The vast majority of them use gradient-based approaches, such as steepest descent, conjugate gradients or Newton–Raphson methods (2), which have the advantage of converging rapidly to the solution closest to the starting point. However, due to the complex profile of the minimized function, it is usually a local minimum, not a global minimum. The necessity to provide an initial guess for the searched parameters is the main drawback of these algorithms since the constants are not known *a priori* and may differ by orders of magnitude. When reasonable preliminary estimates of the kinetic constants are known, deterministic minimization techniques can be used to refine them. However, if preliminary estimates are not available and the character of the newly found minimum is not known, the usual remedy consists of repeating minimization with different starting points and accepting the solution, which is closest to the experimental data. By increasing the number of different starting points, we increase our confidence that the best fit will approach the global minimum (within the limits of a predetermined accuracy). Unfortunately, we can never be certain that the data set cannot be described more satisfactorily by a different combination of parameters. Finally, the overall time necessary to accomplish this task is significantly increased, which greatly reduces the appeal of this approach in a multidimensional case.

Another option is to use stochastic approaches, such as genetic algorithms, simulated annealing or particle swarm optimization, which increase the probability of finding the global minimum (3). There is no guarantee of success, but there is no need for a starting point, and the result depends on adequate sampling of the parameter space, which is user-controlled. Unfortunately, these algorithms are rarely available in widely accessible programs. One has to resort to commercial software, such as MATLAB (4), which may be problematic with regard to the cost and the programming skills it requires.

Another option is to use deterministic global optimization algorithms (5). The problems we need to solve are mixed integer nonlinear programming (MINLP) problems, which represent a severe challenge because few software solutions are available. Most of them are commercial products such as ANTIGONE (6), BARON (7) or Octeract Engine (8), and only a few are open source, e.g., Couenne (9) or SCIP (10). The major drawback is that some programming skills are necessary to effectively use them.

The program presented in this article was born from the frustration of our colleagues, who wanted a simple tool for nonprogrammers, which would reliably analyze their data and help validate multiple models. Consequently, we have developed this program with the following objectives in mind: (i) operation requiring no knowledge of programming techniques, (ii) user-friendly interface for data input and output, (iii) flexibility in defining models for fitting, (iv) customizable graphical visualization of fitting results, (v) comprehensible output of quantitative results, and (vi) cost-free use.

### Implementation

The program’s graphical user interface (GUI) was written in C++, using the Qt library v.5.15.2 (11) with the plotting widget QCustomPlot v2.1.1 (12). For the numerical calculations, a library of Fortran 77 routines was created and interfaced with the GUI. For its development, the Intel oneAPI Base and HPC toolkits (13) were used. The code was compiled and tested on Windows 10, as well as on several Linux platforms: Debian v.11, openSUSE v.15, Ubuntu v.22, Fedora v.38 and Mint v.20.3 (unfortunately, we had no access to the MacOS platform). Two versions are available, either with the installer or without the installer (the so-called “portable” version).

Two algorithms have been implemented: particle swarm optimization (PSO) (14) and simulated annealing (SA) (15). The latter was combined with the conjugate gradient algorithm (16). The setup dialog allows full access to all parameters controlling their behavior. There are five tabs in the GUI that make it possible to do the following:

#### (i) Define the equation, variables, parameters and constants

To define an equation, users can type their own formulae. The parser is flexible and accepts multiline input, temporary variables, and user-defined functions, which can be nested. There are also a number of predefined, intrinsic functions that allow for even greater flexibility. There is also a possibility to work with implicit equations. The current limits on the number of independent variables and parameters are 25 and 100, respectively.

#### (ii) Enter or import experimental data

Experimental data can be entered manually, copied and pasted from external sources, or imported by reading data files (*.dat, *.txt, *.csv…).

#### (iii) Curve fitting with full user control of the relevant parameters

The progress of the fitting can be monitored on the screen. The results can be saved for pairwise model comparisons.

#### (iv) Create a customizable graph of the results and save it in different formats

Users can define symbols for points, line styles for curves and their colors. The title, axis labels and legend items are modifiable. Additionally, custom annotations can be added. The graph can be saved as a PDF file or as an image (svg, png, jpg, tiff, bmp).

#### (v) Create a quantitative output from the fitting and save it in PDF format

The output includes a summary of the input data, target function and numerical results (the optimal parameters, correlation matrix, and quantitative data for model comparisons). The custom graph can also be appended to this report.

The equation parser is flexible enough for the majority of problems to be entered as expressions in the equation field. However, it is not a programming language and does not allow for certain constructs, such as loops. Moreover, it stores equations converted to internal code on a stack, which is interpreted during calculations. While the vast majority of equations encountered in the field of biology can be treated without any problems, we decided to offer an option for more advanced users, who have programming skills, to define equations of arbitrary complexity. Moreover, the compiled functions run faster than the interpreted functions. For this reason, we added the possibility of dynamically attaching an external library (*.dll for Windows, or *.so for Linux) to the program. Then, one can execute one’s own code from the library. The speedup depends on the nature of the problem, but it may easily reach one order of magnitude.

## Results

To compare the performance of our program with that of other software, we ran multiple tests, of which the two most typical ones are reported in this section. The program was executed on a PC equipped with an Intel Core i9 10980XE processor running at 3 GHz, both on Windows and on Linux.

Fig. 1 shows a simple example of a decaying periodic function of time, described by one independent variable and five parameters. The solution is obtained rapidly, in 2 s, independent of the algorithm selected (PSO or SA).

**Fig. 1.**
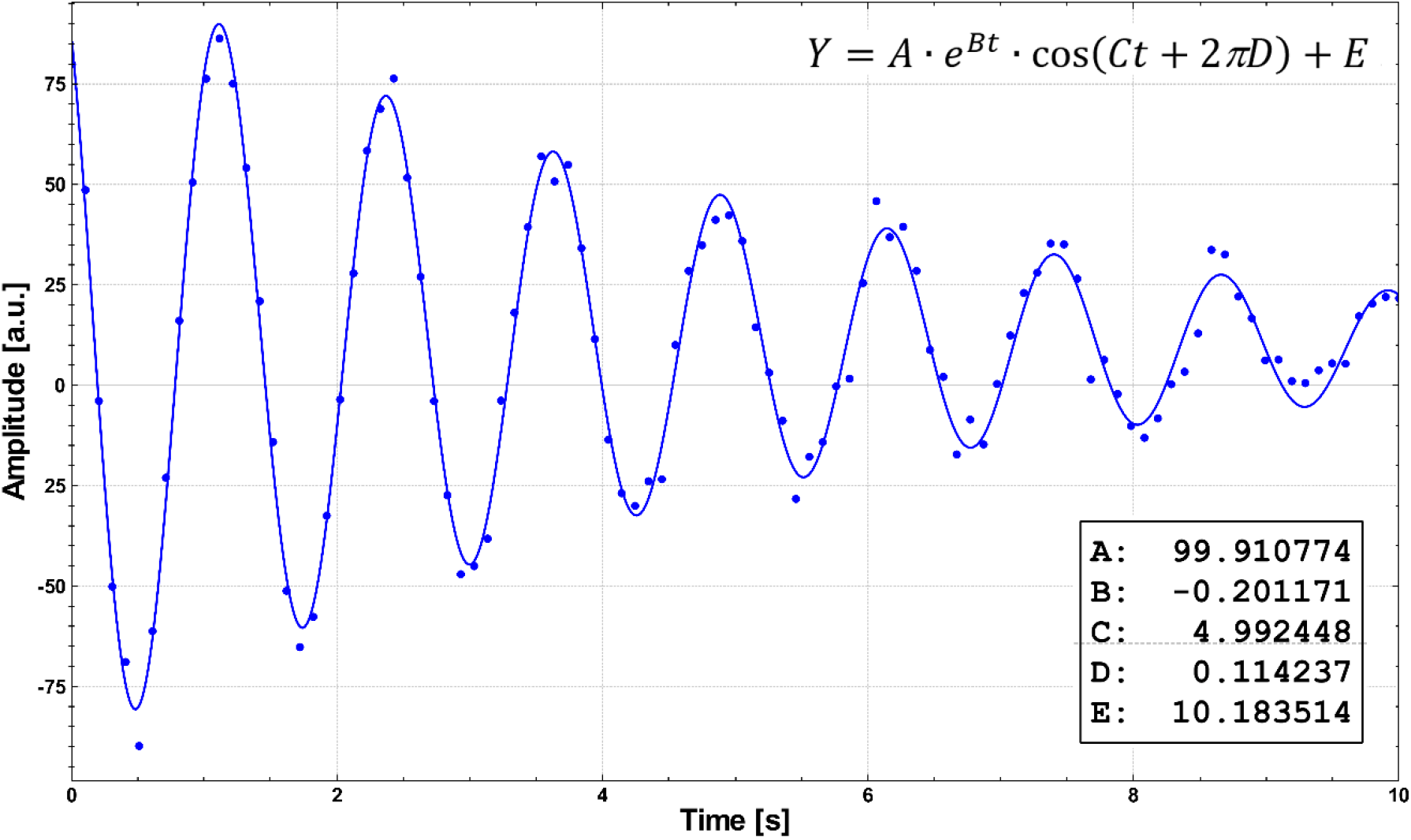
A synthetic data set with random noise representing an NMR free induction decay signal. There is one independent variable (time) and five parameters: A (amplitude), B (decay rate), C (frequency), D (phase) and E (baseline). The curve passing through the points shows the fitting results.

The second test problem we used in evaluating the efficacy of the software concerns a kinetic study of enzymatic reaction pathways. Fig. 2 presents the scheme used in the analysis, and Fig. 3 shows the equations used in the calculations.

**Fig. 2.**
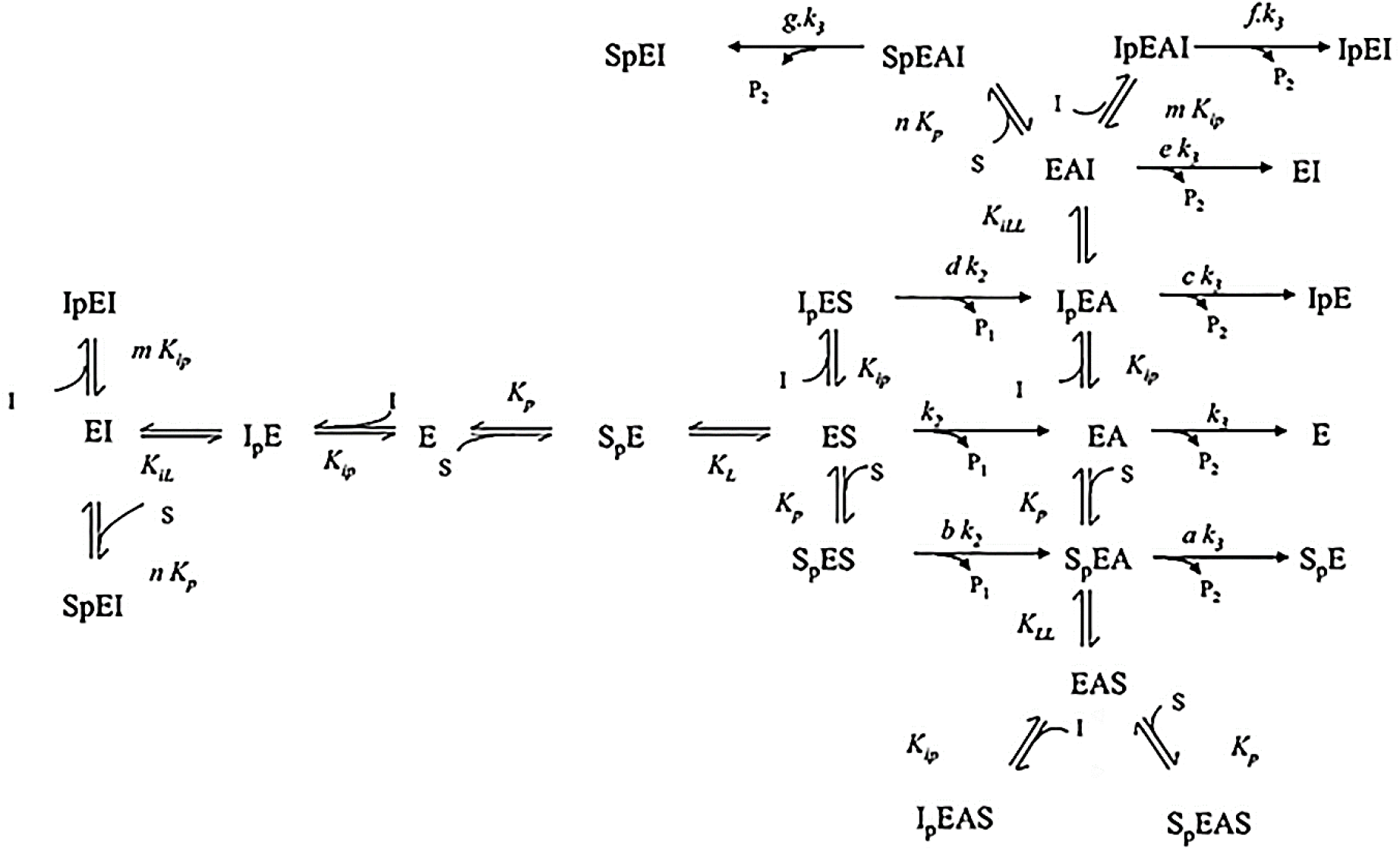
Kinetic study of enzymatic reaction pathways. The example shows the hydrolysis of acetylthiocholine by Drosophila acetylcholinesterase at pH 7 in the presence of edrophonium (courtesy of Prof. Didier Fournier).

**Fig. 3.**
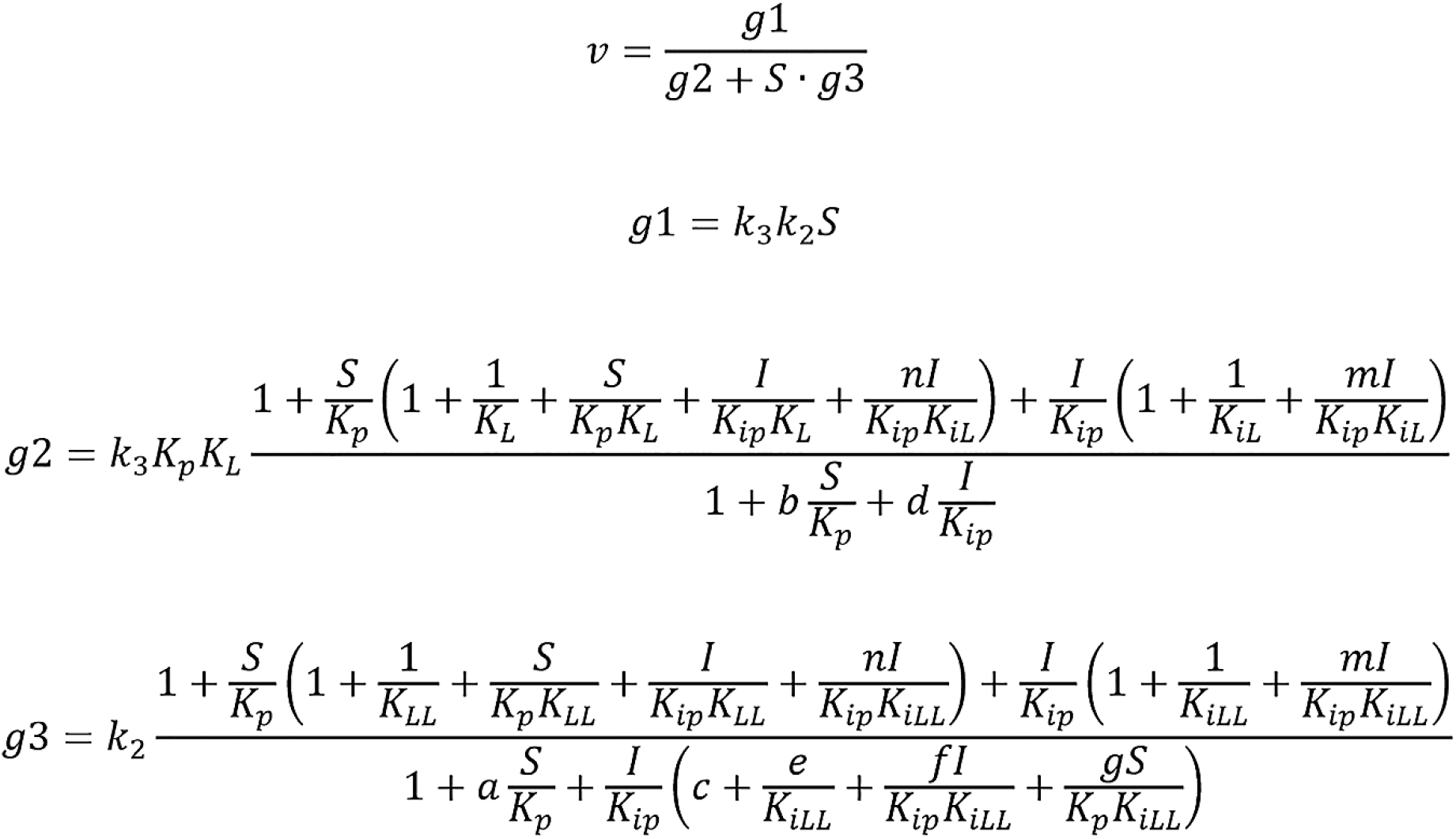
The rate of enzymatic reaction *v* as a function of substrate (S) and inhibitor (I) concentrations. The equations have been derived for the schema presented in Fig. 2.

Although this is not the most complex problem we worked with, it presents a challenge for most programs since it has two independent variables and eleven fitting parameters. We evaluated the performance of the program by measuring the execution times under different conditions. Fig. 4 presents the Glophyt fitting results. The program found the optimum in 23 s using the PSO algorithm and in 151 s using the SA algorithm when using the default settings. It is worth mentioning that when the number of cycles of the SA algorithm (which controls the sampling of the parameter space) is reduced tenfold, the algorithm still yields correct results in 21 s. For comparison, we tested the example from Fig. 2 on identical but precompiled code contained in a dynamically loaded library, and the execution time decreased to 3 s for the PSO algorithm and to 16 s for the SA algorithm (3 s with a reduced number of cycles).

**Fig. 4.**
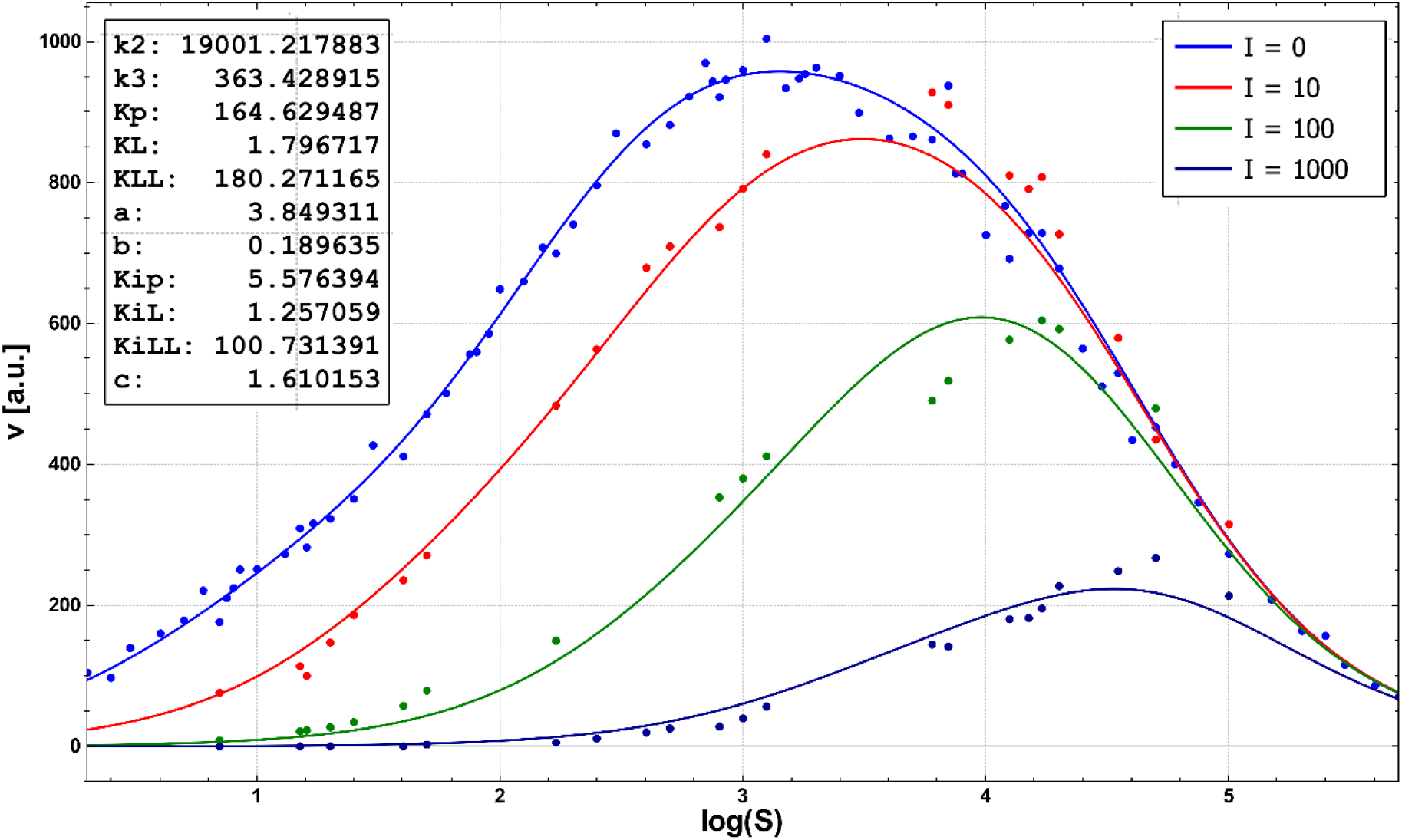
Curve fitting results for the example detailed in Fig. 2. There are two independent variables, the concentrations of substrate (S) and inhibitor (I), as well as fourteen parameters, three of which were fixed (*m = n = 1, e = 0*). The remaining eleven parameters were used in fitting. The values of the fitted parameters span five orders of magnitude, which is sufficient to present a challenge to most of the tested programs.

To compare the performance of Glophyt with those of other programs, we attempted to perform calculations on the test problems using software often cited in the literature. In many cases, a paid license is needed, e.g., for MATLAB (4), OriginLab (17), CurveExpert (18), GraphPad Prism (19) and LAB Fit (20), although sometimes a free trial version is available. There are also free software packages for curve fitting, such as Octave (21), SciDAVis (22), Fityk (23), R/RStudio (24) and Python/SciPy (25). One common feature of these programs is that to solve a least-squares problem, they usually employ gradient-based methods. The fact that they require an initial point is their greatest weakness because a wrong choice leads to a local minimum or even to a lack of convergence.

The first test was conducted on the easy example from Fig. 1. As expected, all tested programs were successful when the selected initial point was close to the optimum. However, when the initial values were randomized or when they were shifted somewhat from the proximity of the minimum, error messages appeared, such as “No convergence” or “Floating point error” (Prism, SciDAVis, Octave, Fityk). In other words, without surprise, we can conclude that gradient-based methods are useful in refining the coordinates of the minimum but not in locating it.

The second example, presented in Fig. 2, was a real challenge. Some of the fitting programs cannot be used because they are limited to one independent variable (Prism, Fityk). Other programs that accept multiple variables have their own limitations, for example, those imposed on the number of parameters to be used in fitting and/or on the length of the equation field, which contains a user-defined function (e.g., 10 parameters and 150 characters for LABFit). The latter is necessary because the function in Fig. 3 cannot be found in libraries often included with the software and needs to be coded as a user-defined function. In our case, we have 11 fitting parameters, and the function requires nearly 300 characters. In general, none of the programs tested in this case yielded positive results (SciDAVis reported “General failure”).

Next, we focused on the software that employs stochastic methods. There are few of them, but the most prominent is MATLAB. We could not test it directly because of the lack of a license, but it was recently demonstrated that the genetic algorithm implemented in MATLAB is not very efficient (26). In the case of Octave, which is free and compatible with MATLAB, the “optim” package seems to be built around gradient-based methods, causing the usual problems with convergence. However, we tend to disregard these software packages for another reason: programming skills are necessary, which goes against one of the basic reasons to create Glophyt. The same is true for R/RStudio and Python/SciPy.

Finally, we tested the deterministic global solvers, which use the branch-and-bound approach, to find the global minimum of a target function by constructing consecutive approximations to its upper and lower limits (27). As before, there are commercial packages (ANTIGONE, BARON, Octeract Engine) and open source software (Couenne, SCIP…). Some of them can handle mixed integer nonlinear programming (MINLP) problems. We tested Couenne within AMPL (28) (free Student version), first on the simple problem mentioned above (one variable, five parameters) and then on the kinetic test problem. In the first case, the program worked as expected, and it displayed correct parameter values. Unfortunately, in the case of the second test problem, the program stopped with an error message and did not find the optimum. However, the major problem is that it is necessary to learn the AMPL language to launch the calculation; hence, it is not meant for nonprogrammers.

To summarize the results, we have not found an alternative for Glophyt within the realm of free and easily accessible software. Our program offers versatility and ease of use that can rarely be found elsewhere. It has already been mentioned before that stochastic algorithms do not guarantee success, but the probability of reaching the optimum is rather high. According to our estimates, based on the ratio of positive results to the overall number of runs, the success ratio is 80-90%. If the result of a given run is not satisfactory, it is advisable to try again because each time the program is launched, it follows a different trajectory through the parameter space; hence, the chances of finding the global minimum increase.

## Conclusions

The objective of this work was to provide researchers with an easy-to-use, reliable program capable of finding global minima of arbitrary functions. The results of test cases conducted on many levels prove that the goal has been achieved. Glophyt succeeds when many other programs fail. It is based on the combination of stochastic and deterministic algorithms and finds the global minimum of a specified function in a given range of variability of unknown parameters in reasonable times (in most cases, from seconds to minutes). For more demanding users, an option is provided to use one’s own precompiled library of equations. The program is available on Windows and Linux platforms, both with an installer and as a portable version, and can be downloaded from the page glophyt.free.fr.

## Availability and requirements

**Project name:** Glophyt

**Project home page:** glophyt.free.fr

**Operating system(s):** Windows, Linux

**Programming language:** C++, Fortran 77

**Other requirements:** None

**License:** GPL3

**Any restrictions to use by non-academics:** None beyond GPL3

## Declarations

### Ethics approval and consent to participate

Not applicable.

### Consent for publication

Not applicable.

### Availability of data and materials

Not applicable.

### Authors’ contributions

GC wrote the program, and SM tested it.

## Acknowledgments

The authors thank Didier Fournier for providing examples of kinetic studies used in testing the program.

